# Fast and Accurate Species Trees from Weighted Internode Distances

**DOI:** 10.1101/2022.05.24.493312

**Authors:** Baqiao Liu, Tandy Warnow

**Author notes:** Corresponding author: Tandy Warnow. Funding: Sandia National Laboratories LDRD. Funding: NSF grant 2006069.

## Abstract

Species tree estimation is a basic step in many biological research projects, but is complicated by the fact that gene trees can differ from the species tree due to processes such as incomplete lineage sorting (ILS), gene duplication and loss (GDL), and horizontal gene transfer (HGT), which can cause different regions within the genome to have different evolutionary histories (i.e., “gene tree heterogeneity”). One approach to estimating species trees in the presence of gene tree heterogeneity resulting from ILS operates by computing trees on each genomic region (i.e., computing “gene trees”) and then using these gene trees to define a matrix of average internode distances, where the internode distance in a tree *T* between two species *x* and *y* is the number of nodes in *T* between the leaves corresponding to *x* and *y*. Given such a matrix, a tree can then be computed using methods such as neighbor joining. Methods such as ASTRID and NJst (which use this basic approach) are provably statistically consistent, very fast (low degree polynomial time) and have had high accuracy under many conditions that makes them competitive with other popular species tree estimation methods. In this study, inspired by the very recent work of weighted ASTRAL, we present weighted ASTRID, a variant of ASTRID that takes the branch uncertainty on the gene trees into account in the internode distance. Our experimental study evaluating weighted ASTRID shows improvements in accuracy compared to the original (unweighted) ASTRID while remaining fast. Moreover, weighted ASTRID shows competitive accuracy against weighted ASTRAL, the state of the art. Thus, this study provides a new and very fast method for species tree estimation that improves upon ASTRID, has comparable accuracy with the state of the art while remaining much faster. Weighted ASTRID is available at https://github.com/RuneBlaze/internode.

## 1 Introduction

Species tree estimation is a common task in phylogenomics and is a prior step in many downstream analyses (e.g., estimating divergence, understanding adaptation). Despite the recent increase in availability of genome-scale data, species tree estimation remains challenging due to gene tree heterogeneity, where gene trees (the evolutionary history of genes) differ from species trees [14]. Among common factors for gene tree heterogeneity, incomplete lineage sorting (ILS), a population-level process modeled statistically by the multi-species coalescent (MSC) [44, 22], is extremely common and well-studied.

A standard approach to species-tree reconstruction under the presence of ILS is to concatenate the alignments of the individual genes and running a maximum likelihood (ML) heuristic on the combined alignment. This simple approach, however, has been established to be statistically inconsistent under the MSC, and can even be positively misleading, converging to the wrong topology with probability 1 as the number of genes increases [40, 39]. Empirically, concatenation can also suffer from degraded accuracy under higher levels of ILS, and can be affected by scalability issues under large data [26, 31]. In response, many accurate ILS-aware methods have since been developed. Those that are most commonly used in practice fall into a class of so-called summary methods, where gene trees are first independently estimated from each genomic region; the inferred gene trees are then used as input to the summary method to “summarize” the input gene trees into a species tree.

In recent years, many accurate summary methods statistically consistent under the MSC have been developed, such as MP-EST [20], NJst [37], ASTRAL [26], ASTRID [46], FASTRAL [8], wQFM [23]. Many of these methods are scalable to genomic-scale data, and under sufficient gene signal and ILS tend to be more accurate than concatenation [31]. Among these methods, ASTRAL is the most commonly used, compares favorably to other methods in accuracy, and has been successfully applied to many large scale data [30, 52]. However, it is well known that summary methods suffer under inaccurate gene trees [31, 49] as a result of summary methods not explicitly taking gene tree uncertainty into account. Under inaccurate gene trees, it might still be preferable to use concatenation even under substantial ILS [31]. Although co-estimating the gene trees and species trees such as using StarBeast [33] or directly inferring quartets from the alignment and then combining them using SVDquartets+PAUP* [5] circumvents this problem, neither can scale to large data.

This sensitivity of summary methods to gene tree error has motivated approaches preprocessing the gene trees to improve the quality of the signal. Although throwing out inaccurate gene trees generally does not help [31], statistical binning [28, 3] and contracting low-support branches [52] improved accuracy for ASTRAL on many conditions. Nonetheless, these approaches require setting arbitrary thresholds: statistical binning requires a threshold to determine which branches are trustworthy, and contracting low-support branches also requires such a threshold. Suboptimal parameter selection in either cases can lead to little accuracy improvement, or even worse, degraded accuracy compared to simply running on the original input [3, 52]. Thus, pragmatically, accurately applying such methods faces the difficulty of parameter selection.

Very recently, Zhang and Mirarab introduced weighted ASTRAL [51]. By directly incorporating gene tree uncertainty into the ASTRAL optimization problem, weighted ASTRAL improved ASTRAL in accuracy under all of their tested conditions. Notably, under conditions where concatenation proved more accurate than unweighted ASTRAL, weighted ASTRAL achieved the largest improvement, shrinking substantially the long known gap [29, 30, 34] between summary methods and concatenation under low gene signal. More specifically, (unweighted) ASTRAL heuristically searches for a species tree that maximizes the amount of quartet trees (unrooted four-taxon tree) shared with the input gene trees. By using branch support and lengths to weigh the reliability of gene tree quartets, weighted ASTRAL instead heuristically maximizes the weighted agreement w.r.t. the input gene trees, effectively discounting the contribution of unreliable quartets. Weighted ASTRAL is threshold-free, was shown to be more accurate than running ASTRAL on contracted gene trees [51], and in fact might be the most accurate summary method under ILS that can scale to large datasets.

Here, inspired by weighted ASTRAL, we introduce weighted ASTRID, incorporating gene tree uncertainty into ASTRID. ASTRID, a fast and more accurate variant of NJst, is based on the internode distance, defined by ASTRID as the number of edges between two taxa in a gene tree. We explore variations of this internode distance where branch uncertainty is considered. Notably, ASTRID is shown to have competitive accuracy against ASTRAL [46, 31] while having a much faster running time [46, 8], both of which we hope to generalize to weighted ASTRID when compared against weighted ASTRAL, obtaining a fast alternative to a very accurate method.

The rest of the study is organized as follows. In Section 2 we describe weighted ASTRID and introduce the two ways of weighting the internode distance matrix before providing some theoretical running time bounds. In Section 3 we describe our experimental study, choosing parameters for weighted ASTRID and comparing it to other methods. In Section 4, we present our experimental results showing that support-weighted ASTRID is very fast, is more accurate than ASTRID, has comparable accuracy with weighted ASTRAL, and provides an alternative accurate species-tree inference method robust to low gene signal, whereas branch-length weighted ASTRID has mixed accuracy and is less accurate than support-weighted ASTRID. In Section 5 we summarize our main conclusions and discuss future work.

## 2 Materials and methods

### 2.1 Basic definitions

Let *n* denote the number of taxa and let *k* denote the number of genes assuming a set of gene trees. Given an unrooted phylogenetic tree *T*, we denote its leafset by *ℒ*(*T*) and its edge-set *E*(*T*). For each edge *e* in *T*, deleting *e* from *T* partitions the leaves into two sets defined by the two connected components separated by *e*. Let this bipartition be denoted by *π*_*e*_ for some *e* ∈ *E*(*T*). We denote the set of bipartitions of *T* by *C*(*T*) = {*π*_*e*_ | *e* ∈ *E*(*T*), and *C*(*T*) uniquely defines the (unrooted) topology of *T*. Two bipartitions are said to be compatible if they can coexist in some tree, and two trees are said to be compatible if their bipartition sets are pairwise compatible. The Robinson-Foulds distance (RF distance) [38] between two trees *T* and *T*′ on the same leafset is the size of the symmetric difference between the bipartitions of *T* and *T*′, i.e., |*C*(*T*) Δ *C*(*T*,′)|. Given two trees *T* and *T*′, we define the nRF (error) rate as their RF distance normalized by | *E*(*T*) | + *E*(*T*′) (the sum of their edge-set sizes), obtaining a value that is between 0 and 1.

For taxa *u, v* ∈ *ℒ* (*T*), let *P*_*T*_ (*u, v*) denote the set of edges on the unique path connecting *u* and *v* in *T*. Given an estimated gene tree *G*, we assume that each internal branch *e* is associated with a branch support value *s*(*e*), some measure of confidence that this branch is correctly estimated, where *s*(*e*) ∈ [0, 1]. Note that we say a branch is correctly estimated (in topology) if the bipartition associated with that edge is present in the true gene tree. We extend this notion of branch support also to pendant edges, where we simply define *s*(*e*) = 1 if *e* is incident to a leaf. Similarly, for a true or estimated gene tree *G*, let *l*(*e*) denote the length of a branch, where for estimated gene trees is usually given in substitution units inferred by some ML tree inference method.

### 2.2 Intertaxon-distance based summary method

Under the context of summary methods, given an input of (unrooted) gene trees, our task is to infer an unrooted species tree topology from these input gene trees. A class of summary methods first introduced by Liu and collaborators [21, 19] infers the species tree by first metricizing the gene trees by defining a specific intertaxon distance between leaf nodes on the gene trees, and then for each pair of taxa averages all such distances across the input gene trees. The final averaged distance, represented as a distance matrix, is then fed as input to a distance-based tree inference method, such as UPGMA [25] or neighbor-joining [41], which infers a tree topology that will be the output species tree of this summary method. Here we focus on ASTRID (a faster and more accurate variant of NJst) and specifically describe its most accurate setting under no missing data.

ASTRID metricizes the gene trees by the **internode distance** *d*_*G*_(*u, v*) where *u, v* ∈ ℒ(*G*) is defined to simply be the number of edges in between *u, v* in *G*, i.e., *d*_*G*_(*u, v*) = |*P*_*G*_(*u, v*)|. Although this definition from ASTRID differs from the original internode distance from NJst [19], where it was defined as the number of nodes (instead of edges) in between two taxa, this change has essentially no impact for our purposes [1] and generalizes to our later modifications better. Given this measure of internode distance, and assuming that each pair of taxa appears at the same time in some gene tree (no missing data), ASTRID proceeds as follows with the input set of gene trees 𝒢:

1. Initialize an *n* × *n* matrix *D* (*n* the number of taxa).
2. For each pair of taxa *u, v*, set *D*[*u, v*] to be the empirical mean of *d*_*G*_(*u, v*) where *G* ranges in the input gene trees 𝒢 where both *u, v* are present, i.e., set 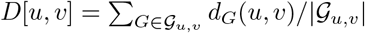, where 𝒢_*u,v*_ is {*G* | *G* ∈ 𝒢 ∧ *u, v* ∈ *ℒ*(*G*)}. This empirical mean is well defined as we assumed |*𝒢*_*u,v*_| *>* 0.
3. Run FastME’s heuristic for balanced minimum evolution [15] (referred to as FastME from here on) on *D*, outputting an unrooted species tree.

ASTRID is statistically consistent under the MSC when given true gene trees [1], and is in practice very fast while having competitive accuracy [46, 31].

### 2.3 Weighted ASTRID

In the definition of the internode distance, each branch contributes equally to the internode distance for each pair of taxa it separates. Intuitively, under the realistic assumption that gene trees are estimated with non-trivial amount of error, some branches will be more reliably estimated than the others. As such it makes sense to assign weights to branches as some confidence of them correctly contributing to the internode distance. For example, because zero-support branches very likely do not contribute correctly to the internode distance, assigning a weight of 0 to such branches likely discounts incorrect contributions to the internode distance. The branch lengths could also be used as such a proxy, because short branches are empirically hard to estimate. Thus, our problem becomes to choose appropriate weighting schemes for the edges based on information already annotated in the gene trees, that is, the branch support and branch lengths. Because branch support is already designed as some statistical confidence of the correctness of some branch, it seems natural to naively assign the support directly as the weight for each branch. We alternatively explore simply assigning the branch length as the weight. The details are presented as follows.

#### 2.3.1 Distance defined by branch support

We now formally introduce **wASTRID-s** (weighted ASTRID by support), analogous to the naming of weighted ASTRAL by support. As mentioned above, if a branch has low support, it is less likely that this branch contributes correctly to the internode distance, and thus branches with low support should contribute less. Here we try one simple approach, defining each branch’s contribution to the internode distance as its support instead of 1, which gives rise to the following definition of *d*_*G*_(*u, v*), the new support-weighted intertaxon distance replacing the internode distance from step 2 of ASTRID:

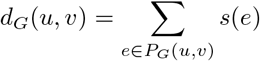

In reality, several different measures of support exist with different running time and accuracy trade-offs [2]. Weighted ASTRAL discovered that the approximate Bayesian support [2] of IQ-TREE led to the most accurate species tree reconstruction, although other measures of support also led to accuracy improvements over unweighted ASTRAL. We leave this choice of support as a parameter to be decided later for wASTRID-s.

While ASTRID is statistically consistent under the MSC when given true gene trees [1], the statistical consistency of wASTRID-s only makes sense under *estimated* gene trees because only estimated gene trees can have meaningful branch support. We conjecture that similar to wASTRAL-s (support weighted ASTRAL), wASTRID-s is statistically consistent under the MSC under some probablistic interpretation of branch support when given estimated gene trees.

#### 2.3.2 Distance defined by branch lengths

Unlike branch support, which is designed to be a measure of statistical confidence on the correctness of a branch, branch lengths can only serve as proxies to such information, where shorter branches are empirically harder to estimate likely as a result of shorter branches containing less information (fewer substitutions) [47]. We do not attempt a complex conversion here, and simply just assign the branch length as the confidence similar to how we use the support values:

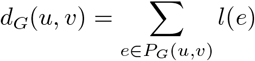

Notably, this definition of *d*_*G*_(*u, v*) coincides with STEAC [21] (motivated from a different perspective of the coalescence time between genes), and potentially under a more accurate setting when paired with the FastME step of ASTRID. In addition, we also explore whether and how to normalize the input branch lengths of the gene trees for the weighting. We name this final algorithm **wASTRID-pl** (weighted ASTRID by path-lengths).

### 2.4 Running time and fast distance calculation

Clearly, both wASTRID-s and wASTRID-pl retain the original theoretical running time of ASTRID. Recall that *n* is the number of species and *k* is the number of gene trees given in the input. For each gene tree, our metricization simply assigns already-computed values as lengths to each edge, thus calculating the intertaxon distance across all pairs of taxa per gene tree takes *O*(*n*^2^) time (for normalizing branch lengths in wASTRID-pl, we only explore ways to normalize that does not affect this asymptotic running time). The averaged intertaxon distance thus can be calculated in *O*(*kn*^2^) time. Under the reasonable assumption of the Yule-Harding distribution [50, 10] on the output species trees, the balanced minimum evolution heuristic of FastME takes in expectation *O*(*n*^2^ lg *n*) time [7]. Therefore we re-derive the original running time result:

#### Theorem 1.

*The averaged intertaxon distance of wASTRID-s and wASTRID-pl can be obtained in O*(*kn*^2^) *time, with the final species tree obtained using an additional O*(*n*^2^ lg *n*) *time by FastME assuming the output species tree is under the Yule-Harding distribution, where n is the number of species and k is the number of genes*.

While the algorithm for the *O*(*kn*^2^) step can be theoretically uninteresting, we implemented another algorithm for the distance calculation in hope of better in-practice speed. The asymptotically optimal algorithm is easy to devise because the naive algorithm, which given a gene tree, starts a BFS at each leaf to obtain the all-pairs intertaxon distance, is already quadratic time per gene and also asymptotically optimal due to each gene tree having 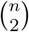 distances. The original ASTRID implementation, in this vein, uses an algorithm which implicitly performs multiple traversals in the tree. For weighted ASTRID, we instead implemented an intertaxon-distance algorithm from TreeSwift [32] based on post-order traversal, through which we hope to achieve better empirical performance due to better cache locality in its simultaneous maintenance of multiple distances from the leaves in an array.

## 3 Experimental Study

### 3.1 Overview

We conduct three experiments. In Experiment 1, we explore parameter choices (choice of branch support for wASTRID-s and normalization scheme for wASTRID-pl) for weighted ASTRID. In Experiment 2, we compare the accuracy and running time of weighted ASTRID against other methods on a diverse set of simulated conditions. In Experiment 3, we compare ASTRID, wASTRAL-h, and the best variant of wASTRID as determined by previous experiments on the Jarvis et al. [12] avian biological dataset comparing the quality of the reconstructed species trees.

### 3.2 Datasets

We assembled a set of diverse data from prior studies (see Table 1), consisting of various simulated conditions with estimated gene trees and one biological dataset (“avian biological”) from the avian phylogenomics project [12]. We use the nomenclature of the original ASTRID study and refer to the SimPhy-simulated datasets from the ASTRAL-II study by a “MC” name. The ILS levels of the datasets are measured in average discordance (AD), defined as the average nRF rate between the true species tree and the true gene trees. Same as the original ASTRID study [46], we classify the ILS levels of the datasets into four categories according to their AD percentages, where below 25% is classified as low ILS (L), between 26% and 39% mid ILS (M), between 40% and 59% high ILS (H), and higher AD considered very high ILS (VH). For the simulated conditions, we subsample the gene trees to size *k* = 50, 200, 1000 except for the mammalian simulation (*k* = 50, 100, 200 instead).

**Table 1:** Dataset statistics. The ILS levels of the datasets are categorized according to their AD percentages, where below 25% is low ILS (L), between 26% and 39% mid ILS (M), between 40% and 59% high ILS (H), and higher AD very high ILS (VH). SH-like denotes FastTree default support; BS denotes standard bootstrap support using FastTree or RAxML; aBayes denotes IQ-TREE approximate Bayesian support. (*): training dataset: only 10/50 replicates were used for training.

| Dataset | # taxa | # genes | # reps | ILS (AD %) | Branch Support |
| --- | --- | --- | --- | --- | --- |
| ASTRAL-II MC2* [30] | 201 | 1000 | 10 | 33 (M) | SH-like, BS, aBayes |
| ASTRAL-III S100 [52] | 101 | 1000 | 50 | 46 (H) | aBayes |
| ASTRAL-II MC3 [30] | 201 | 1000 | 50 | 21 (L) | aBayes |
| ASTRAL-II MC5 [30] | 201 | 1000 | 50 | 34 (M) | aBayes |
| ASTRAL-II MC1 [30] | 201 | 1000 | 50 | 69 (VH) | aBayes |
| ASTRAL-II MC6H [18] | 201 | 1000 | 50 | 9 (L) | aBayes |
| ASTRAL-II MC11H [18] | 1001 | 1000 | 50 | 35 (M) | aBayes |
| Avian 2x [28] | 48 | 1000 | 20 | 29 (M) | aBayes |
| Avian 1x [28] | 48 | 1000 | 20 | 47 (H) | aBayes |
| Avian 0.5x [28] | 48 | 1000 | 20 | 60 (VH) | aBayes |
| Mammalian 2x [28] | 37 | 200 | 20 | 21 (L) | aBayes |
| Mammalian 1x [28] | 37 | 200 | 20 | 29 (M) | aBayes |
| Mammalian 0.5x [28] | 37 | 200 | 20 | 50 (H) | aBayes |
| Jarvis et al. avian [12] | 48 | 14446 | 1 | not known | BS |

For the measure of support for the datasets, the weighted ASTRAL study provided gene trees reannotated with aBayes support and IQ-TREE inferred branch lengths for the ASTRAL-II and ASTRAL-III datasets. We also reannotated the avian and mammalian simulation with aBayes support and IQ-TREE branch lengths because aBayes was determined as the best measure of support also for wASTRID-s. We use the MC2 condition as the training data for both wASTRID-s and wASTRID-pl, where to explore the choice of branch support for wASTRID-s, we also took the original FastTree [35] inferred trees annotated with the default FastTree SH-like support and also computed a version of the gene trees with standard bootstrap support [9] using 100 FastTree trees.

### 3.3 Methods

For testing, we compare against ASTRID, ASTRAL, and weighted ASTRAL, with the following settings:

- **ASTRID** (v2.2.1), the base method of weighted ASTRID. We use the original implementation, turning off missing data imputation, as our data has no missing entries in the final averaged matrix.
- **ASTRAL(-III)** (v5.7.8). We do not contract very low support edges in the gene trees because a good threshold can be hard to determine across datasets. In addition, no contraction allows us the fully explore the impact of weighting.
- **wASTRAL-h** (hybrid weighted ASTRAL, v.1.4.2.3). This was the most accurate weighted ASTRAL from the original study, using both branch lengths and support to weigh gene tree quartets. wASTRAL-h supports parallelization, so we run wASTRAL-h with 16 threads.

Many of the datasets have FastTree-inferred gene trees that were reannotated with IQ-TREE approximate Bayesian support. FastTree-inferred trees have polytomies for identical sequences, but these polytomies will be resolved when the trees get reannotated by IQ-TREE, adding false positive edges which may adversely affect the accuracy of the unweighted methods. In these cases, we run the unweighted methods on the original FastTree gene trees.

### 3.4 Evaluation criterion

For simulated datasets, we compare the topological error rate of the reconstructed species trees using the normalized Robinson-Foulds error (nRF error) w.r.t. the true species trees. Because all the inferred and true species trees are binary, the nRF error rate is equivalent to the missing branch rate (ratio of branches in the reference tree missing from the reconstructed tree).

On the avian biological dataset, as the true tree is not known, we compare the estimated species trees against prior topologies (wASTRAL tree and published trees). We also compute the local posterior-probability (localPP) branch support [42] for the the reconstructed species trees obtained using wASTARL-h to assess the reliability of the branches.

On all datasets, we keep track of the wall-clock running time of the methods, the time taken from consuming the input gene trees (that may have been preprocessed with new branch support values) until outputting the species tree.

### 3.5 Experimental Environment

All experiments were conducted on the Illionis Campus Cluster, a heterogeneous cluster that has a four-hour running time limit. The heterogeneity of the hardware makes the wall-clock running times not directly comparable across runs, but can still be used to gather obvious running time trends.

## 4 Results & Discussion

### 4.1 Experiment 1: Parameter selection

After exploring the choice of branch support among the default FastTree SH-like support, IQ-TREE approximate Bayesian (aBayes) support (normalized to the [0, 1] range), and bootstrap support (100 FastTree replicates) on the training dataset (figure omitted, shown in Appendix Figure 7), the best accuracy of wASTRID-s is obtained by using the normalized aBayes support on gene trees. All measures of support, however, improved the species tree estimation error in general. The superiority of (normalized) aBayes support is consistent with the support chosen for weighted ASTRAL, where it was also found superior to SH-like support and bootstrap support. This advantage is even more pronounced when considering that aBayes support can be obtained much faster than bootstrap support [2].

On this training dataset, wASTRID-pl attained the highest accuracy when normalizing the branch lengths in each gene tree by the maximum path length in that gene tree (better than no normalization). Interestingly, while worse than ASTRID with fewer genes *k* ∈ {50, 200}, wASTRID-pl attained higher accuracy than ASTRID when *k* = 1000. However, when comparing wASTRID-s and wASTRID-pl, wASTRID-pl was always less accurate.

### 4.2 Experiment 2: Results on simulated datasets

In this experiment, we show four-way comparisons among ASTRID, ASTRAL, weighted ASTRAL (wASTRAL-h), and weighted ASTRID (wASTRID-s). We omit showing the branch-length weighted wASTRID-pl, as it was discovered to be on all datasets less accurate than wASTRID-s. We put an emphasis on the accuracy (nRF error), while later revisiting the problem of running time.

#### 4.2.1 ASTRAL-III S100

This 101-species dataset contains four conditions that varied in the gene tree estimation error (GTEE, measured by the average nRF error between the estimated gene trees and their corresponding true gene trees) by varying the sequence lengths. We show the results of three of the four conditions in Figure 1. In all such figures, we show the unweighted methods (ASTRID, ASTRAL) in dotted lines. On S100, aside from the obvious trend that summary methods become more accurate as *k* (the number of genes) increases, all methods also improve in accuracy when given more accurate estimated gene trees. These two trends are unsurprising and well-documented across studies for summary methods in general.

**Figure 1:**
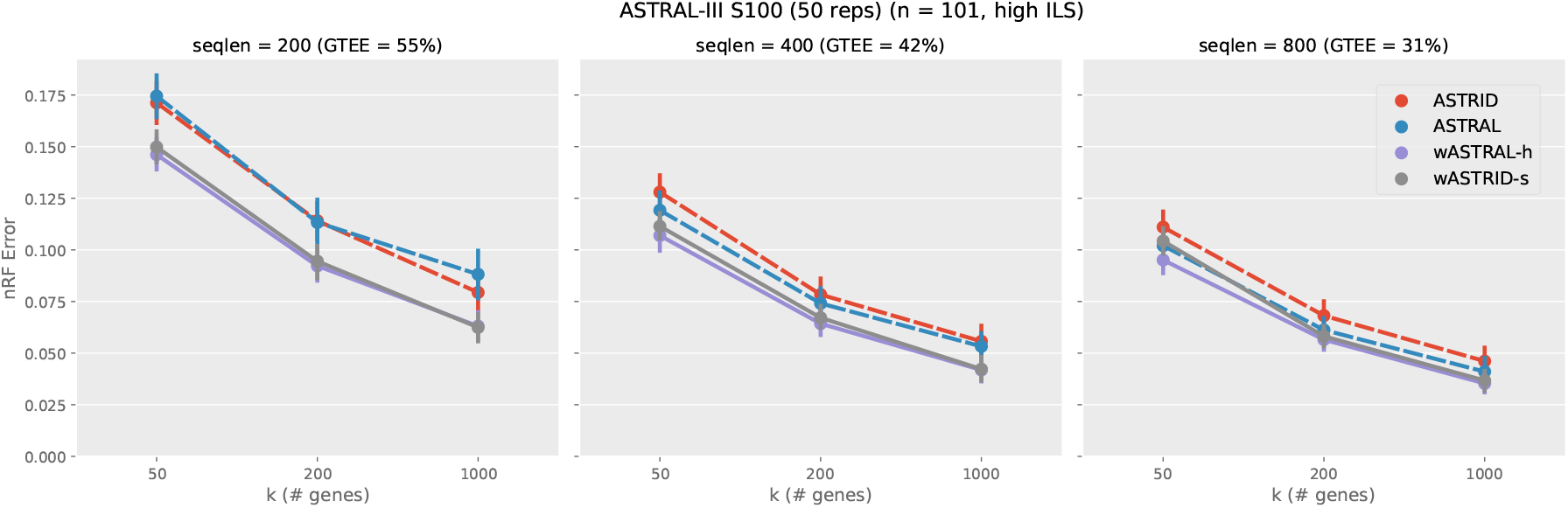
Topological error of species tree across methods on the ASTRAL-III S100 dataset (*n* = 101, AD = 46%). Subfigures vary the sequence lengths, affecting the gene tree estimation error (measured in GTEE, the average distance between estimated gene trees and true gene trees). The *x*-axis varies in the number of genes. Results are shown averaged across 50 replicates with standard error bars. All weighted methods (wASTRID, wASTRAL) ran on gene trees reannotated with IQ-TREE aBayes support branch support and lengths. All methods achieves better accuracy when given more (larger *k*) or better (lower GTEE) data. Weighted methods are more accurate than unweighted ones. wASTRID-s and wASTRAL-h have almost the same accuracy.

**Figure 2:**
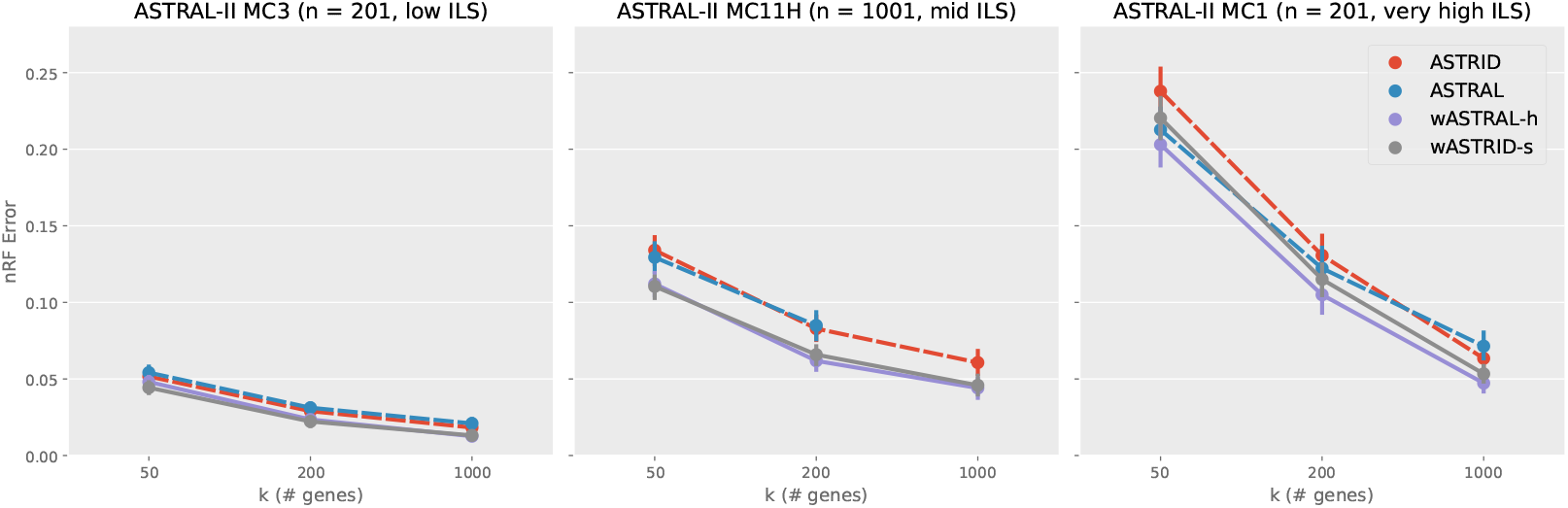
Topological error (nRF error rate) of species tree across methods on selected conditions on the ASTRAL-II SimPhy conditions (*n* = 201, 1001, 201, AD = 21, 35, 69% respectively). Each subfigure depicts a different model condition. The *x*-axis varies in the number of genes. Results are shown averaged across 50 replicates with standard error bars. ASTRAL did not finish 24 out of the 50 replicates within four hours for *k* = 1000 on MC11H and thus the data point was omitted. All weighted methods (wASTRID, wASTRAL) were run on gene trees reannotated with IQ-TREE aBayes support branch support and lengths. Weighted methods are more accurate than unweighted ones. wASTRID-s on MC1 was less accurate than wASTRAL-h and otherwise has the same accuracy.

More interestingly, the weighted methods (wASTRID-s, wASTRAL-h) are clearly more accurate than their unweighted counterparts, especially at higher levels of GTEE (GTEE = 0.55, 0.42). The improvement in accuracy from the weighted methods does not seem to depend on the number of genes, suggesting that the noise brought by low quality gene trees is not simply resolved by having ample data. This advantage of the weighted methods, however, is smaller as more accurate gene trees are used (GTEE = 0.31), as expected.

wASTRID-s clearly improves upon ASTRID on this dataset, with an obvious advantage in the nRF error across all conditions and all number of genes. wASTRID-s notably almost matches the accuracy of wASTRAL-h. With *k* ∈ {200, 1000}, no clear benefit exists for using wASTRAL-h on the shown conditions.

On this dataset, wASTRID-pl (as seen in full results in Appendix Figure 8), similar to trends seen in the training dataset, attains better accuracy than ASTRID when *k* = 1000, but is almost always worse than wASTRID-s. While this better accuracy than ASTRID suggests potential for wASTRID-pl and STEAC-like methods in general, this accuracy disadvantage to wASTRID-s led to us dropping wASTRID-pl from future experiments.

#### 4.2.2 ASTRAL-II SimPhy

This dataset was generated varying the speciation rates, ILS level, and number of taxa, with the “H”-suffixed conditions regenerated from a previous study [18] halving sequence lengths to increase GTEE. We show the results in 2 (only three out of five of the conditions visualized for brevity. See Appendix Figure 10 for the full figures).

Across all conditions in this dataset, the weighted methods are more accurate than the unweighted methods. This advantage does not seem to depend on the level of ILS or the number of species. Even under the easiest condition (MC3), wASTRAL-h and wASTRID-s still consistently achieved better accuracy. All methods also performed worse in accuracy as ILS increases, as expected.

While wASTRID-s still consistently improved upon ASTRID in accuracy on this dataset, we also see datasets where wASTRID-s is worse than wASTRAL-h (MC1 as shown here and MC6H as shown in Appendix Figure 10). The relative performance of wASTRID-s and wASTRAL-h seems related to the relative performance of the base methods: MC1 and MC6H are the two conditions that ASTRAL is in general more accurate than ASTRID, but the relative performance of the base methods does not explain the whole picture – for MC1 going to *k* = 1000, ASTRID became more accurate than ASTRAL yet wASTRID-s is still worse than wASTRAL-h. More positively, on the other conditions of this dataset, wASTRAL-h has nearly the same accuracy as wASTRID-s, although wASTRAL-h is marginally more accurate, which might be due to the hybrid weighting of wASTRAL-h also using the branch lengths.

On the largest input of these conditions (MC11H, *k* = 1000), ASTRAL did not finish under our four-hour time limit on around half of the replicates (see Appendix Section B.1 for more details), but wASTRAL-h did. We comment on this scalability advantage of wASTRAL-h over ASTRAL later.

#### 4.2.3 Avian and mammalian simulation

These two datasets were generated based on model trees inferred on biological datasets. Both datasets have three conditions with varying ILS by scaling the model tree branch lengths by 2X, 1X, or 0.5X, with shorter branch lengths leading to higher degrees of ILS. Notably, prior results from the ASTRID study [46] showed that ASTRID outperformed ASTRAL on the avian simulation, while on the mammalian simulation ASTRAL was more accurate. Also, the mammalian simulation only has 200 genes available, so we vary *k* among 50, 100, 200 unlike the other datasets.

On the avian simulation (Figure 3), aside from obvious trends (ILS increases difficulty; more genes leads to more accurate reconstruction), same as the original study, ASTRID is consistently more accurate than ASTRAL. Strangely, although the weighted methods inherit the relative performance of their base methods, in a few cases the weighted methods do not help in accuracy, but they do not erode accuracy either. Even on conditions where the weighted methods improved accuracy, the improvement was small. For example, wASTRAL-h, even though improving upon ASTRAL, is even less accurate than ASTRID, whereas on previously shown data wASTRAL-h was consistently the best in accuracy. This avian simulation does carry substantial GTEE (*>* 50%), so it is not clear what led to the weighted methods underperforming.

**Figure 3:**
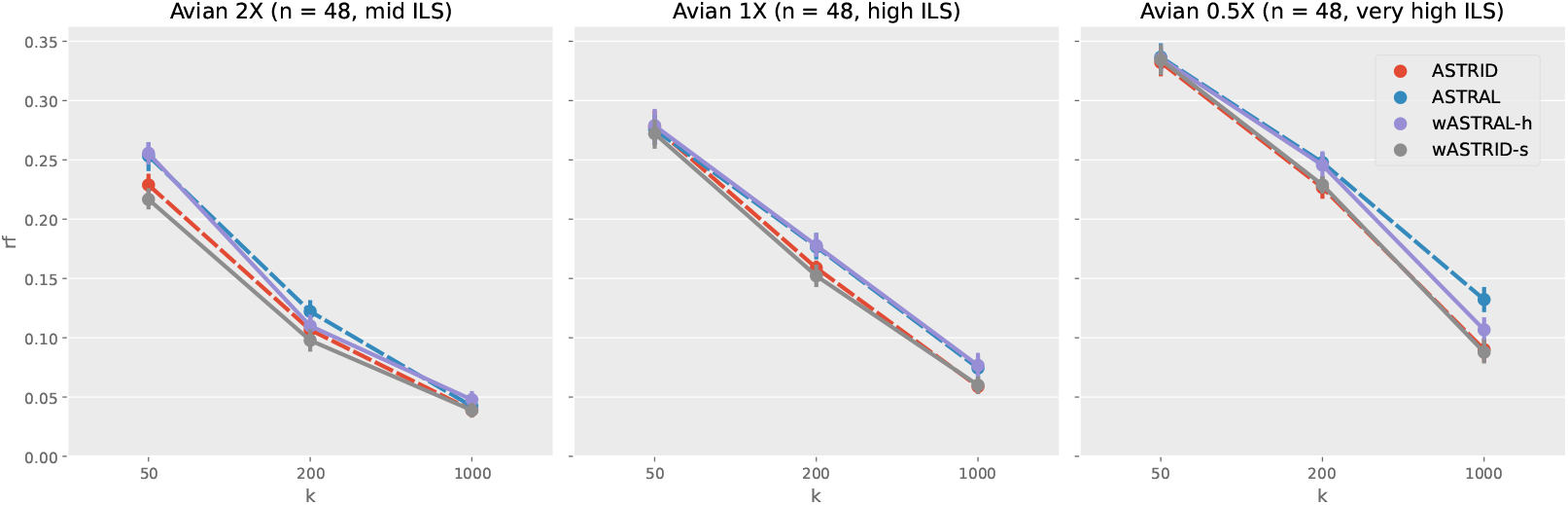
Topological error (nRF error rate) of species tree across methods on the avian simulation (*n* = 48). Each subfigure depicts a different model condition. The *x*-axis varies in the number of genes. Results are shown averaged across 20 replicates with standard error bars. All weighted methods (wASTRID, wASTRAL) ran on gene trees reannotated with IQ-TREE aBayes support branch support and lengths. ASTRID and wASTRID-s is more accurate than ASTRAL and wASTRAL-h, with a slight accuracy advantage to the weighted methods over the unweighted ones.

The results for the mammalian simulation (Figure 4) paints a more perplexing picture. On the 2X condition, surprisingly, the weighted methods are less accurate than their unweighted counterparts in general. This trend continues with the 1X condition, where wASTRAL-h only mostly matches ASTRAL in accuracy, and wASTRID-s is worse than ASTRID in accuracy. Only at the 0.5X condition, both weighted methods clearly help in accuracy. wASTRAL-h is clearly better than wASTRID-s on this dataset, but this difference can be explained by the accuracy advantage of ASTRAL on ASTRID. While it is again unclear why the weighted methods underperformed, this dataset is relatively easy compared to the previously shown datasets, with all methods achieving around 0.05 nRF error rate even with *k* = 200, so despite the puzzling relative performance, the difference in accuracy among methods is very small.

**Figure 4:**
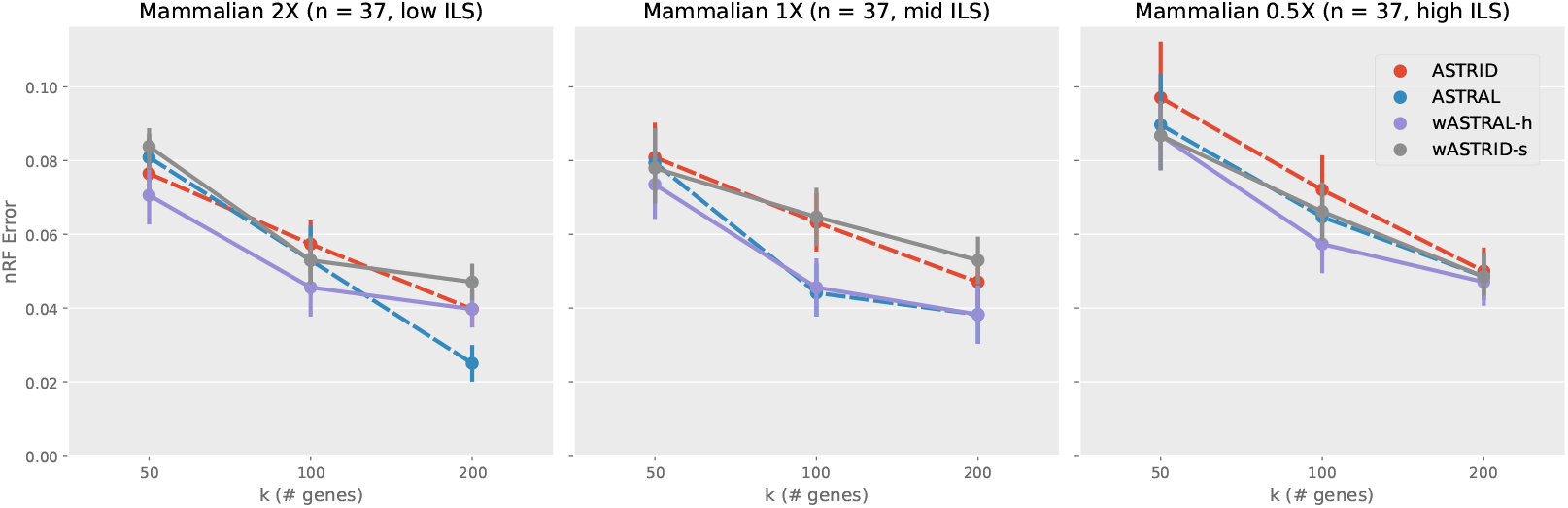
Topological error (nRF error rate) of species tree across methods on the mammalian simulation (*n* = 37). Each subfigure depicts a different model condition. The *x*-axis varies in the number of genes. Results are shown averaged across 20 replicates with standard error bars. All weighted methods (wASTRID, wASTRAL) ran on gene trees reannotated with IQ-TREE aBayes support branch support and lengths. ASTRAL and wASTRAL-h is more accurate than ASTRID and wASTRID-s. The weighted methods have mixed accuracy compared to the unweighted ones.

#### 4.2.4. Running time

We show the wall-clock running time of the four methods under three representative conditions (*n* = 101, 201, 1001) in Table 2, with a direct comparison of the two most accurate methods visualized in Figure 5. While the heterogeneity of the hardware dilutes the comparability of the running times, clearly wASTRID-s and ASTRID are much faster than wASTRAL-h and ASTRAL, with wASTRID-s on average taking less than 12 seconds even on the largest input, whereas on the same input wASTRAL-h on average takes roughly two hours. In general, wASTRID-s is around two magnitudes faster than wASTRAL-h. Although we note that the default flags of both ASTRAL and wASTRAL-h (that we used in the experiments) also calculate support and lengths for reconstructed species tree, in practice this is a fast step relative to the species tree reconstruction, and does not affect our running time analysis in any substantial way. On MC11H, wASTRID-s and ASTRID took less time going from *k* = 50 to 200, likely due to the *k* = 50 trees having larger diameters, negatively impacting the FastME step running time which has a linear dependency on the diameter of the output tree.

**Table 2:** Wall-clock running time (sec) across methods on selected representative simulated conditions on *n* = 101, 201, 1001 for *k* ranging in 50, 200, 1000. Data points show averages across 50 replicates. ASTRAL did not finish 24 out of the 50 replicates within four hours for *k* = 1000 on MC11H and thus the data point was omitted. The methods sorted by the fastest to the slowest is almost always wASTRID-s, ASTRID, wASTRAL-h, and ASTRAL across all shown conditions. ASTRID and wASTRID-s are much faster than ASTRAL and wASTRAL-h.

| Running time (s) | k (# genes) | ASTRAL | ASTRID | wASTRAL-h | wASTRID-s |
| --- | --- | --- | --- | --- | --- |
| S100 ($n = 101$) | 50 | 12.1 | 0.1 | 4.6 | 0.1 |
|  | 200 | 29.6 | 0.2 | 6.6 | 0.1 |
|  | 1000 | 300.7 | 1.4 | 22.8 | 0.2 |
| MC5 ($n = 201$) | 50 | 20.3 | 0.2 | 12.3 | 0.1 |
|  | 200 | 57.5 | 0.6 | 25.6 | 0.2 |
|  | 1000 | 600.7 | 1.8 | 106.9 | 0.5 |
| MC11H ($n = 1001$) | 50 | 552.9 | 18.0 | 611.8 | 11.3 |
|  | 200 | 2059.6 | 14.9 | 1333.4 | 8.8 |
|  | 1000 | – | 27.5 | 7191.0 | 11.8 |

**Figure 5:**
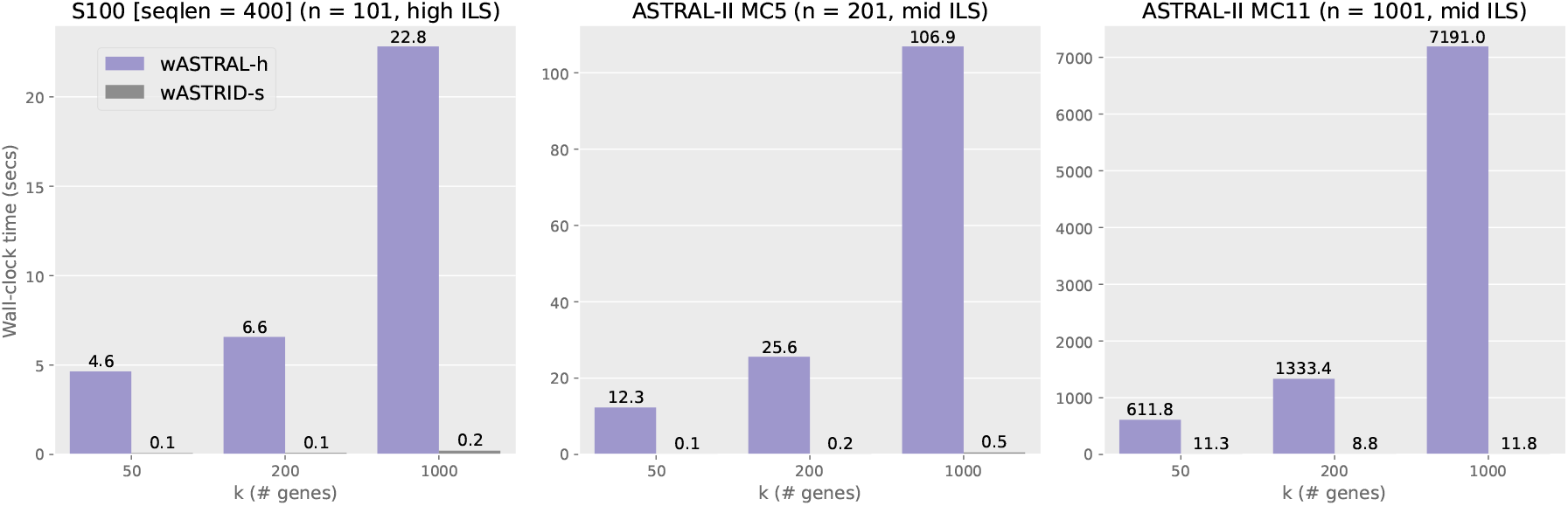
Wall-clock running time (sec) comparison of wASTRID-s and wASTRAL-h on selected representative simulated conditions on *n* = 101, 201, 1001 for *k* ranging in 50, 200, 1000. Bars and labels show averages across 50 replicates. wASTRID-s is dramatically faster than wASTRAL-h.

The weighted methods are by large faster than their unweighted counterparts. For example on S100 (seqlen = 400), wASTRAL-h under *k* = 1000 is more than ten times faster than ASTRAL, and ASTRAL did not finish around half of the datasets for the largest input (MC11H, *k* = 1000). Aside from the benefit of parallelization (we ran wASTRAL-h using 16 threads, but off-the-shelf ASTRAL does not support parallelization), this speed advantage under large number of genes of wASTRAL-h over ASTRAL can also be attributed to the algorithmic change implemented in wASTRAL-h. The new weighted ASTRAL algorithm removes the in-practice quadratic dependency of ASTRAL’s search algorithm on the number of genes [27] and instead has a linear dependency on *k* for the running time. wASTRID-s is consistently faster (at least two times faster on most conditions) than ASTRID, showing that the new distance calculation algorithm implemented is more efficient than the original one.

### 4.3 Experiment 3: Results on the avian biological dataset

Jarvis et al. studied the phylogeny of birds using a dataset on 48 taxa using 14446 genes [12]. The original gene trees were annotated with RAxML [43] bootstrap support, which we directly use in our wASTRID-s analysis. This dataset is known to have very low gene resolution, with the average support only 32% [28]. ASTRAL on the original set of gene trees reconstructed a species tree that contained obvious inaccuracies, but contracting low support branches enabled ASTRAL to construct a very plausible topology in agreement with established results [52]. Zhang and Mirarab reanalyzed the original gene trees using wASTRAL-h, and reconstructed the same topology as ASTRAL running on contracted gene trees [51]. In addition to directly comparing against their wASTRAL-h tree, we also compute the number of differing branches between the inferred trees and the two published trees of the Jarvis et al. study: the ExaML-based concatenation tree, called the TENT (“total evidence nucleotide tree”), and the coalescent based published tree based on binned MP-EST (MP-EST* tree). We show the results in Figure 6. In addition to showing the topology and the localPP branch support, we also highlight in gradients six clades that are thought to be strongly corroborated [4] for avian phylogeny.

**Figure 6:**
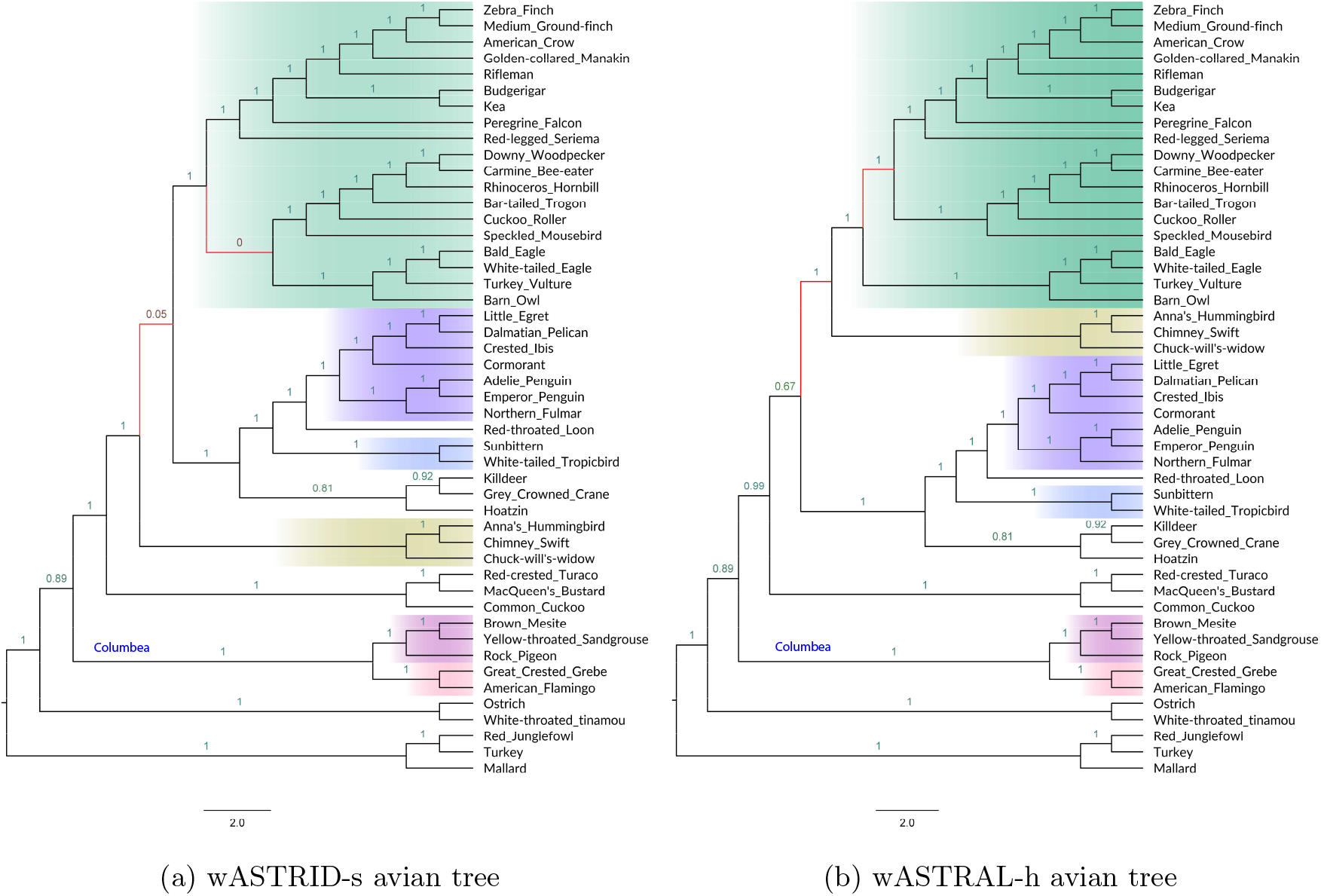
Results on the avian biological dataset (*n* = 48, *k* = 14446). In (a) and (b), we show the reconstructed species tree topology of wASTRID-s and wASTRAL-h, annotated with local posterior-probability branch support (localPP) computed using wASTRAL-h. The red branches show where the two trees differ. The six well corroborated clades according to Braun and Kimball [4] are displayed by both trees and highlighted in gradients. For wASTRID-s, the red branches also coincide with the only two low support branches. Contracting the very low support branches for wASTRID-s arrives at a topology compatible with wASTRAL-h. wASTRAL-h took around 294 seconds to infer its tree. wASTRID-s was very fast and took 1 second to infer its topology.

The ASTRID tree (shown in Appendix Figure 9) is relatively discordant with the wASTRAL-h tree and the reference trees. Compared to the wASTRAL-h tree it differed in 8 branches, and compared to the TENT and MP-EST* tree it differed in 9 and 8 branches. The ASTRID tree also did not recover the Columbea clade, a clade that has seen strong support in various analyses of this data [12, 52, 36].

Using wASTRID-s, we recovered a topology that is much more in agreement with the other trees, differing in 2 branches (4.4% of the branches) with the wASTRAL-h tree. It is also in much higher agreement with the published trees (differing in 3 branches with the TENT, 2 branches with the MP-EST* tree). Looking closer, the two branches where it differs from the wASTRAL-h tree coincide with the only two very low support branches (no more than 5% in support), and thus contracting the very low support branches arrives at a topology compatible with the wASTRAL-h tree. Both wASTRID-s and wASTRAL-h recover the Columbea-Passerea split, a major conclusion in the original analysis of this data, and even agree on the placement of the hard-to-place Hoatzin.

For running time, both ASTRID and wASTRID-s finished quickly. ASTRID completed in 6.4 seconds, and wASTRID-s completed in 1.1 seconds. Our rerun of wASTRAL-h on this data finished in 294.2 seconds, showcasing the much better scalability of the new weighted ASTRAL optimization algorithm in the number of genes, whereas ASTRAL on the same input took 32 hours in the ASTRAL-III study. In summary, on this dataset, wASTRID-s inferred a more accurate tree compared to ASTRID, is much faster than wASTRAL-h, and is compatible with wASTRAL-h after contraction of two very low support branches.

### 4.4 Additional remarks

#### Impact of number of genes, ILS, and GTEE

In all results shown, all methods achieved better accuracy when given more genes, and all methods have decreased accuracy as the degree of ILS or GTEE increases. These trends are well known and quite expected for all summary methods. The number of species does not seem to have an influence on the accuracy of the methods. The relative performance of the methods does shift across different numbers of genes, but there seems to be no reliable predictor of this change across the conditions. All methods seem equally adversely impacted by ILS in accuracy, but for GTEE, the weighted methods are seen to be more robust to the low signal, as by design the weighted methods can extract better signals from the gene trees under high GTEE.

#### Impact of weighting

Weighting improved accuracy on all the datasets (simulated, biological) except on the mammalian simulation, and the improvement of accuracy tends to be the same degree for both wASTRAL-h and wASTRID-s, with a slight advantage to wASTRAL-h, likely due to the hybrid weighting using more information than support-weighted ASTRID. Even with ample genes or lower levels of GTEE, the weighted methods still in general improved upon the unweighted variants in accuracy. It is not clear how the mammalian simulation (and to some degree, the avian simulation) differs from the other datasets, where the weighted methods could in cases be worse than the original methods in accuracy. The mammalian simulation has a non-trivial amount of GTEE, but has the smallest number of species among the tested datasets and was generated from a different protocol than the ASTRAL simulation datasets, and all these factors could potentially affect the accuracy. Although the weighted methods are in general faster than their unweighted counterparts, this speed advantage is orthogonal to the weighting and entirely due to faster algorithmic design or implementation.

#### wASTRAL-h vs. wASTRID-s

ASTRAL and ASTRID are known to be among the most accurate summary methods under ILS, and their relative accuracy is dataset dependent, as also shown in our results. The relative accuracy of their weighted counterparts is clearly influenced by the accuracy of their base methods, as seen in the performance differences in the avian and mammalian simulation, where either wASTRID-s and wASTRAL-h can be the more accurate method. On the ASTRAL simulated datasets (ASTRAL-III S100, ASTRAL-II SimPhy), wASTRID-s and wASTRAL-h have comparable accuracy, with a small advantage to wASTRAL-h, which might be due to wASTRAL-h’s more effective hybrid weighting. Somewhat unsurprisingly, on all conditions except the mammalian simulation, wASTRID-s is more accurate than unweighted ASTRAL, even on conditions where ASTRAL is more accurate than ASTRID.

As for running time, wASTRID-s is clearly the winner. Despite wASTRAL-h’s improved algorithm and parallelism, wASTRID-s is still two magnitudes faster than wASTRAL-h, and hence can provide better scalability on large scale data. We do not extend this running time discussion to a highly parallelized setting (e.g., 64 cores, common for large-scale data analysis) since wASTRAL-h has not been efficiently parallelized yet (as noted in their future work), so any current discussion likely does not reflect the future parallel efficiency of wASTRAL-h. The wASTRID-s algorithm is trivially parallelizable in its distance calculation (albeit very fast already), but parallelizing the FastME step can be difficult both in design and in implementation, which can be a bottleneck under large *n*. Intriguingly, wASTRAL-h theoretically can scale better in the number of species than wASTRID-s as wASTRAL-h has a running time that is subquadratic in the number of species, but our results show that at *n* = 1001 this point is far from being reached. Thus wASTRID-s is much faster than wASTRAL-h, while having comparable accuracy.

## 5 Conclusions

While the estimation of species trees using summary methods, such as ASTRAL, ASTRID, MP-EST, and others, is now commonplace, it is known that gene tree estimation error reduces the accuracy of the estimated species tree. We presented support weighted ASTRID (wASTRID-s), an improvement over ASTRID that incorporates uncertainty in gene tree branches into its estimation of the averaged internode distance. Our work is largely inspired by the recent work of weighted ASTRAL (wASTRAL-h), which improved upon ASTRAL, and we showed that wASTRID-s obtained similar accuracy improvements over ASTRID. The advantage provided by wASTRID-s over ASTRID is most noteworthy under higher degrees of gene tree estimation error. wASTRID-s has very close accuracy to wASTRAL-h and is sometimes more accurate, but overall wASTRAL-h has a small advantage in accuracy. The improvement of wASTRAL-h over wASTRID-s may be due to its weighting also incorporating branch lengths. A branch-length weighted version of ASTRID (wASTRID-pl) has mixed accuracy compared to ASTRID, and does not compete against the support-weighted wASTRID-s in accuracy. In general, wASTRID-s and wASTRAL-h serve as accurate species tree inference methods under ILS and are more robust to GTEE than ASTRID and ASTRAL, two of the most accurate summary methods. Both can scale very well, with wASTRID-s much faster. However, the differences in accuracy are dataset dependent, just as the comparison between ASTRAL and ASTRID for accuracy seems dataset dependent.

This study was limited to datasets where the only cause for gene tree discordance with the species tree was ILS and gene tree estimation error. When considering real world datasets that may have other sources of gene tree heterogeneity, such as GDL or HGT, it seems likely that ASTRAL and other quartet-based methods may have an advantage over ASTRID and wASTRID-s, due to the theoretical proofs of statistical consistency for quartet-based methods for those conditions (and the lack of such proofs for distance-based species tree estimation methods) [16, 24, 6, 11].

Although wASTRID-s is highly accurate and very fast, rather than suggesting that wASTRID-s replace wASTRAL-h as the recommended species tree estimation method, we recommend using wASTRID-s in conjunction with wASTRAL-h and other species tree estimation methods. Due to its speed, the inclusion of wASTRID-s adds little computational burden, and having multiple different approaches for estimating the species tree, each based on a very different (but statistically consistent) technique, can provide insights into what parts of the species tree are most reliably recovered, and which parts may need further data in order for full resolution.

For future work, finding a way to incorporate branch lengths into the branch certainty scoring for wASTRID-s, a hybrid weighting, will improve accuracy and might close the accuracy gap between wASTRID-s and wASTRAL-h under some conditions. Missing data in the final averaged distance matrix has not been addressed, and generalizing the approach from ASTRID [45] for handling missing entries to weighted ASTRID will be important. ASTRID has been used to speed-up and sometimes improve analyses for ASTRAL by assisting in constraining the search space explored by ASTRAL (e.g,, FASTRAL [8]), suggesting that a weighted version of ASTRID might provide even better accuracy improvements in FASTRAL. Another direction for future work is species tree estimation in the presence of gene duplication and loss, where gene family trees have multiple copies of species and so are called MUL-trees. The combination of DISCO [48], a method for decomposing the MUL-trees into single-copy gene trees, with ASTRID produced very good accuracy and scalability [48], suggesting that combinnig DISCO with wASTRID might be even more accurate.

## A Commands

### A.1 Weighted ASTRID

The commands differ depending on the type of support annotated on the gene trees. For FastTree SH-like, bootstrap support, and aBayes support, the commands for wASTRID-s are respectively:

~~~
wastrid -i $genes -o $output # FastTree SH-like
wastrid -x 100 -i $genes -o $output # bootstrap
wastrid -n 0.333 -i $genes -o $output # aBayes
~~~

For the final setting of wASTRID-pl (distance defined by path lengths, with branch lengths normalized by the maximum path-length in tree), the above commands need to be appended with an additional flag: `-m n-length` (distance using “normalized branch length” instead of the default `-m` support).

### A.2 Other summary methods

#### A.2.1 Weighted ASTRAL

We ran hybrid-weighted ASTRAL (v1.4.2.3) using the following command on trees annotated with aBayes support:

~~~
astral-hybrid -x 1 -n 0.333 -i $genes -o $output
~~~

#### A.2.2 ASTRID

We ran ASTRID (v2.2.1) using the following command:

~~~
ASTRID -s -i $genes -o $output
~~~

The `-s` flag is specified to skip the missing data imputation step of ASTRID, as all tested data have no missing data in the averaged internode distance matrix (i.e., each pair of taxa appears together at least once in some gene tree).

#### A.2.3 ASTRAL

We ran ASTRAL (v5.7.8) using the following command:

~~~
java -jar astral.5.7.8.jar -i $genes -o $output
~~~

### A.3 Obtaining gene tree support

#### A.3.1 Approximate Bayesian support

We computed IQ-TREE (v2.1.2) aBayes support using the following command:

~~~
iqtree2 -s $aln -te $gtree -m GTR+G -abayes
~~~

where `$aln` is the alignment file for the gene tree, and `$gtree` is the path to the gene tree topology.

#### A.3.2 Bootstrap support

We computed bootstrap support (on the training dataset) using FastTree (v2.1.11) on bootstrap replicates generated by Goalign (v0.3.5) [17].

The following command was used to generate bootstrap replicates (`$aln` is the original gene alignment):

~~~
goalign build seqboot -i $aln -S -n 100
~~~

The FastTree command used for inferring a tree from one bootstrap replicate is (`$bs aln` is one bootstrap alignment replicate generated by Goalign):

~~~
FastTree -nt -gtr -nosupport $bs_aln > $output
~~~

The FastTree bootstrap trees are then randomly resolved to eliminate polytomies, and then mapped by RAxML-NG [13] to the original gene trees using the following command:

~~~
raxml-ng --support --tree $gtree --bs-trees $bs_trees --bs-metric fbp
~~~

where $gtree is the path to the original gene tree, and $bs trees contains new-line separated newick trees inferred on the bootstrap replicates.

## B Supplementary Text

### B.1 Failures to complete

ASTRAL failed to complete 24/50 replicates (replicates 2, 4, 6, 8, 9, 11, 13, 15, 16, 17, 20, 23, 27, 28,

29, 30, 32, 36, 38, 39, 40, 41, 48, and 49) on the MC11H condition (*n* = 1001) under *k* = 1000. The 24 log files indicate that ASTRAL has indeed timed out on each of the replicates. An example of the last three lines of the log files is attached:

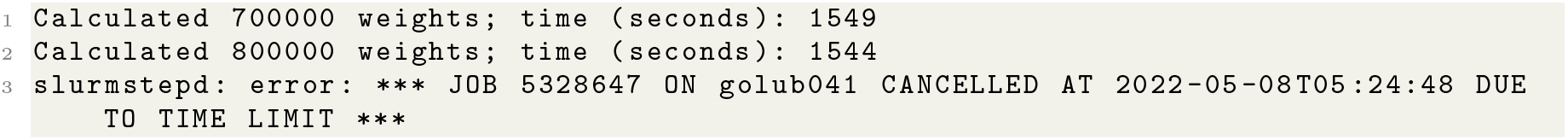

## C Supplementary Figures & Tables

**Figure 7:**
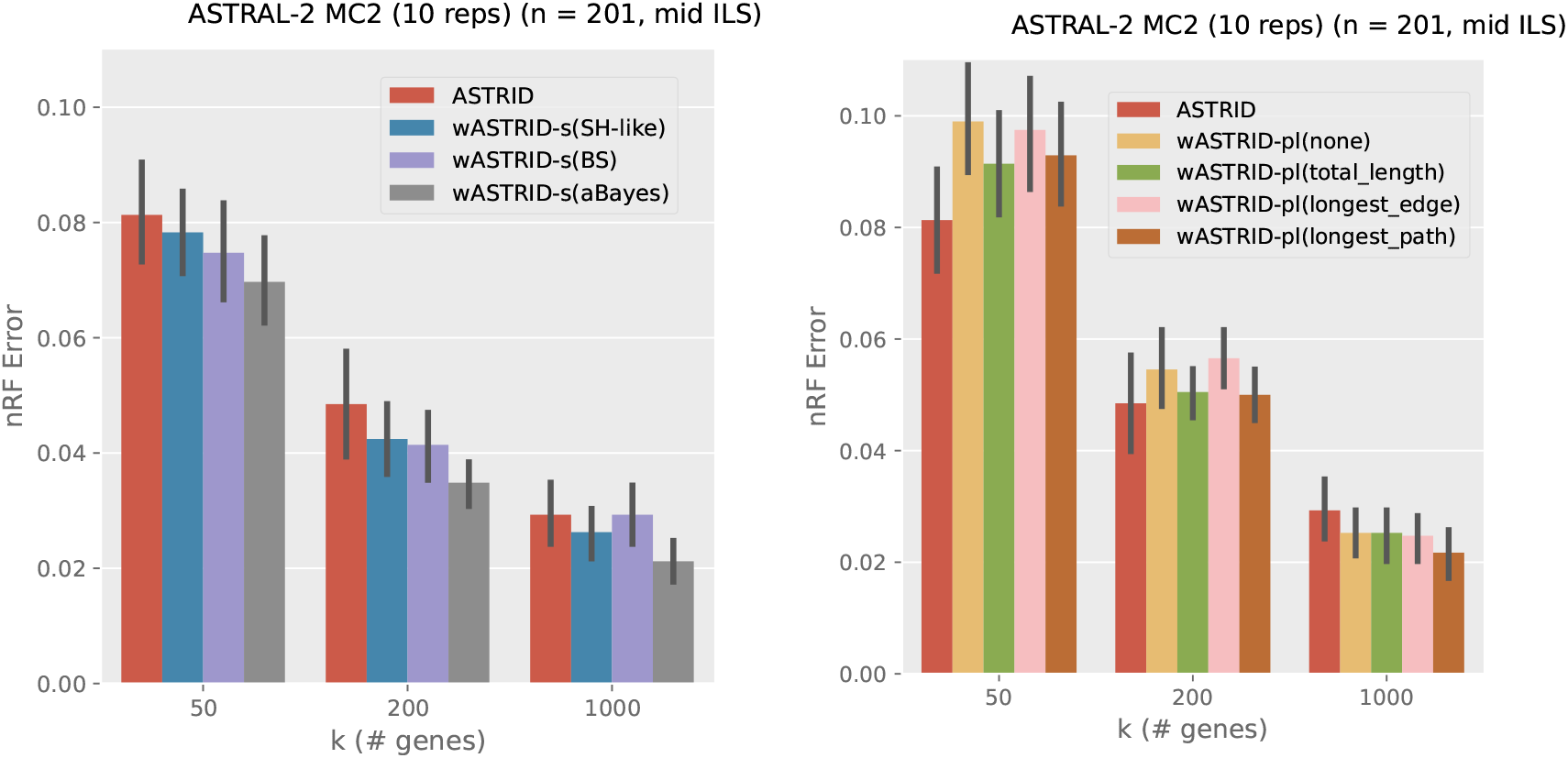
Comparison of the choice of branch support and the choice of branch length normalization strategy for wASTRID-s and wASTRID-pl respectively on the training data, showing the species tree topological error rates (nRF error). “SH-like” is FastTree SH-like support. “aBayes” is IQ-TREE approximate Bayesian support. “BS” is bootstrap support using 100 FastTree trees. The *x*-axis varies in the number of genes *k* in the input. Results are shown averaged across ten replicates. Error bars show standard error.

**Figure 8:**
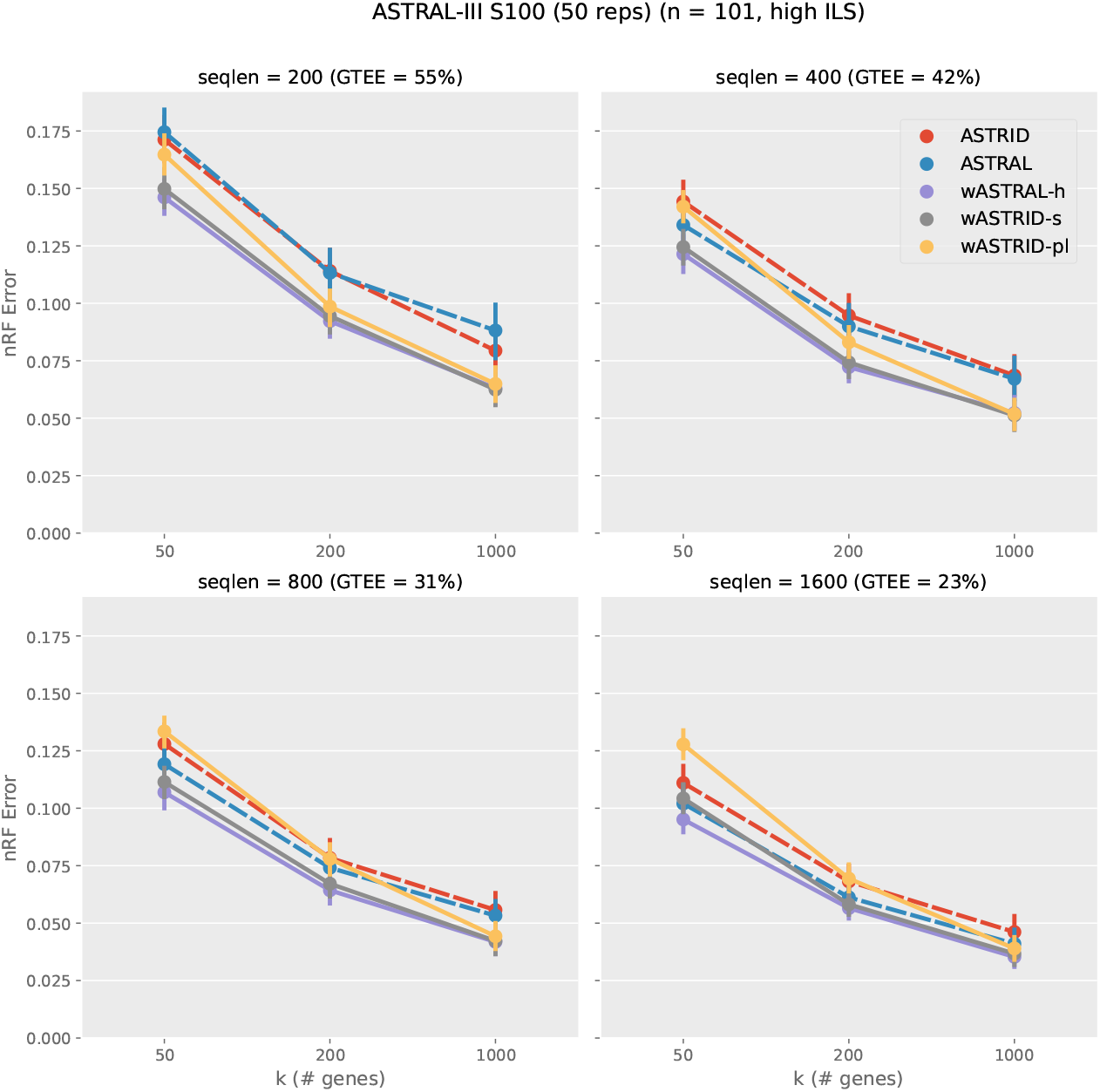
Topological error of species tree across methods on the ASTRAL-III S100 dataset (*n* = 101, AD = 46). Subfigures vary the sequence lengths, affecting the gene tree estimation error (measured in GTEE, the average distance between estimated gene trees and true gene trees). The *x*-axis varies in the number of genes. Results are shown averaged across 50 replicates, with error bars showing the standard error. All weighted methods (wASTRID-s, wASTRID-pl, wASTRAL) ran on gene trees reannotated with IQ-TREE aBayes support branch support and lengths.

**Figure 9:**
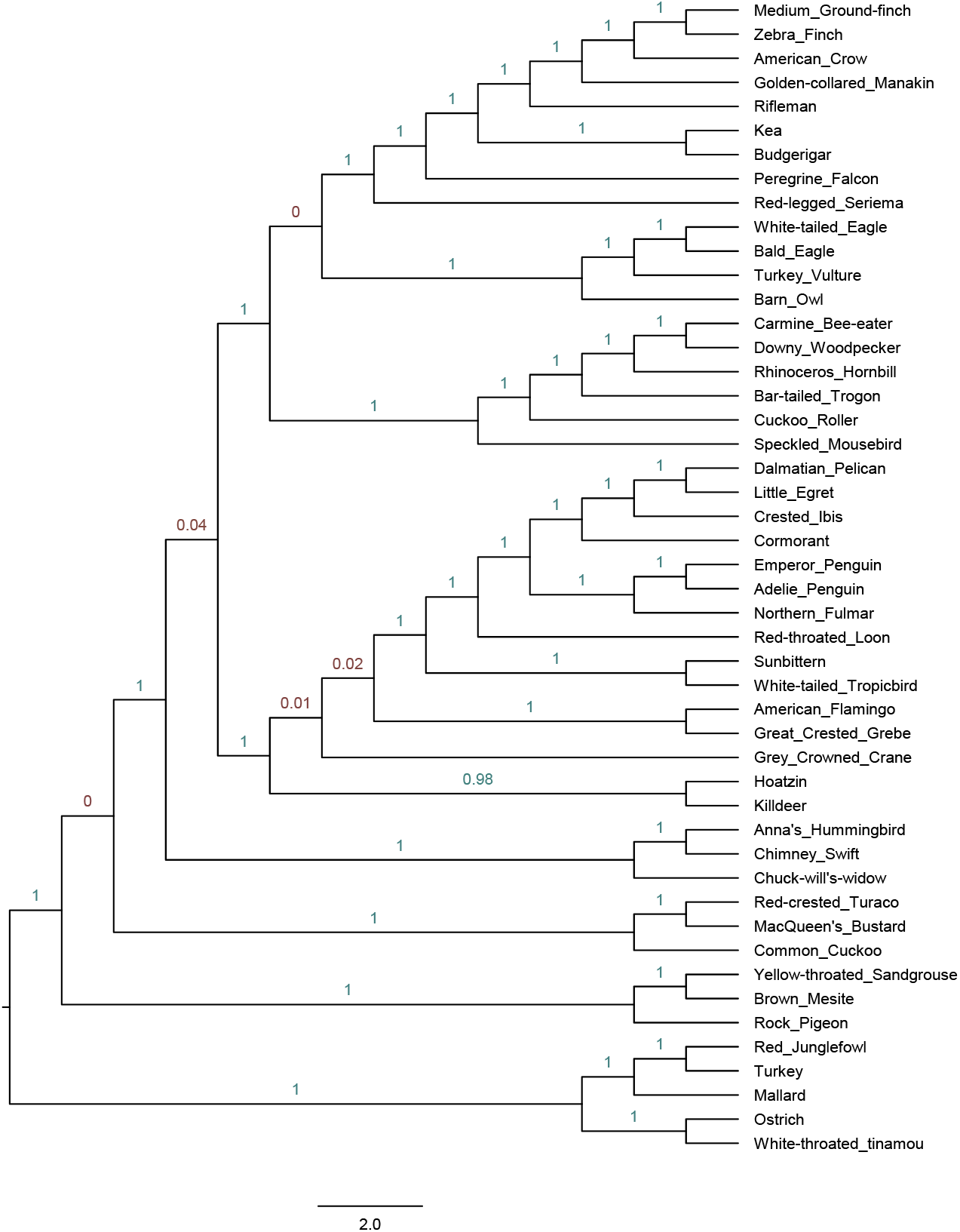
Reconstructed species tree on the Jarvis et al. avian biological data using ASTRID. Branch support are in localPP values calculated by wASTRAL-h.

**Figure 10:**
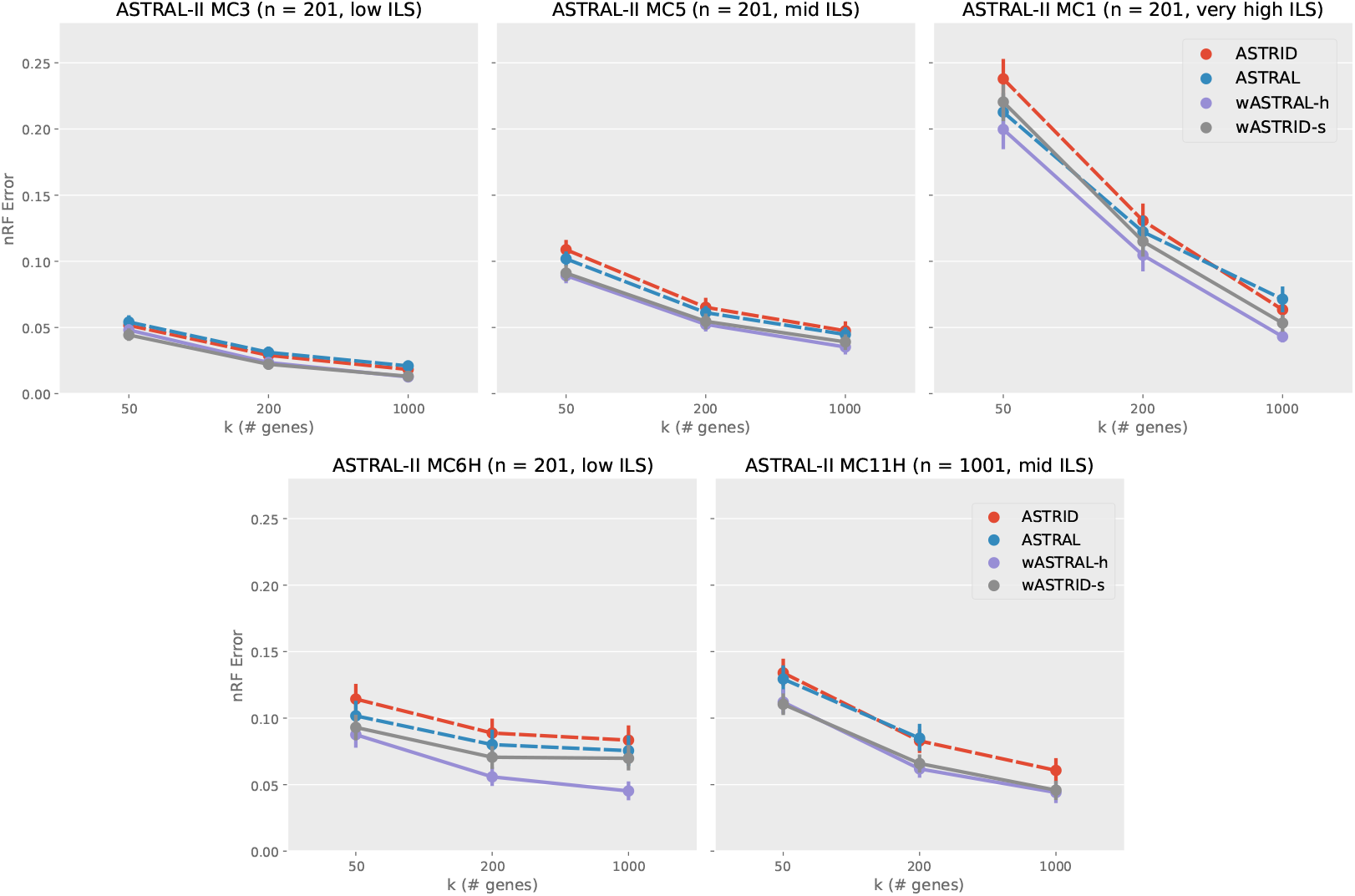
Topological error (nRF error rate) of species tree across methods on selected conditions on the ASTRAL-II SimPhy conditions. Each subfigure depicts a different model condition. The *x*-axis varies in the number of genes. Results are shown averaged across 50 replicates with standard error bars. ASTRAL did not finish 24 out of the 50 replicates within four hours for *k* = 1000 on MC11H and thus the data point was omitted. All weighted methods (wASTRID, wASTRAL) were run on gene trees reannotated with IQ-TREE aBayes support branch support and lengths.

**Figure 11:**
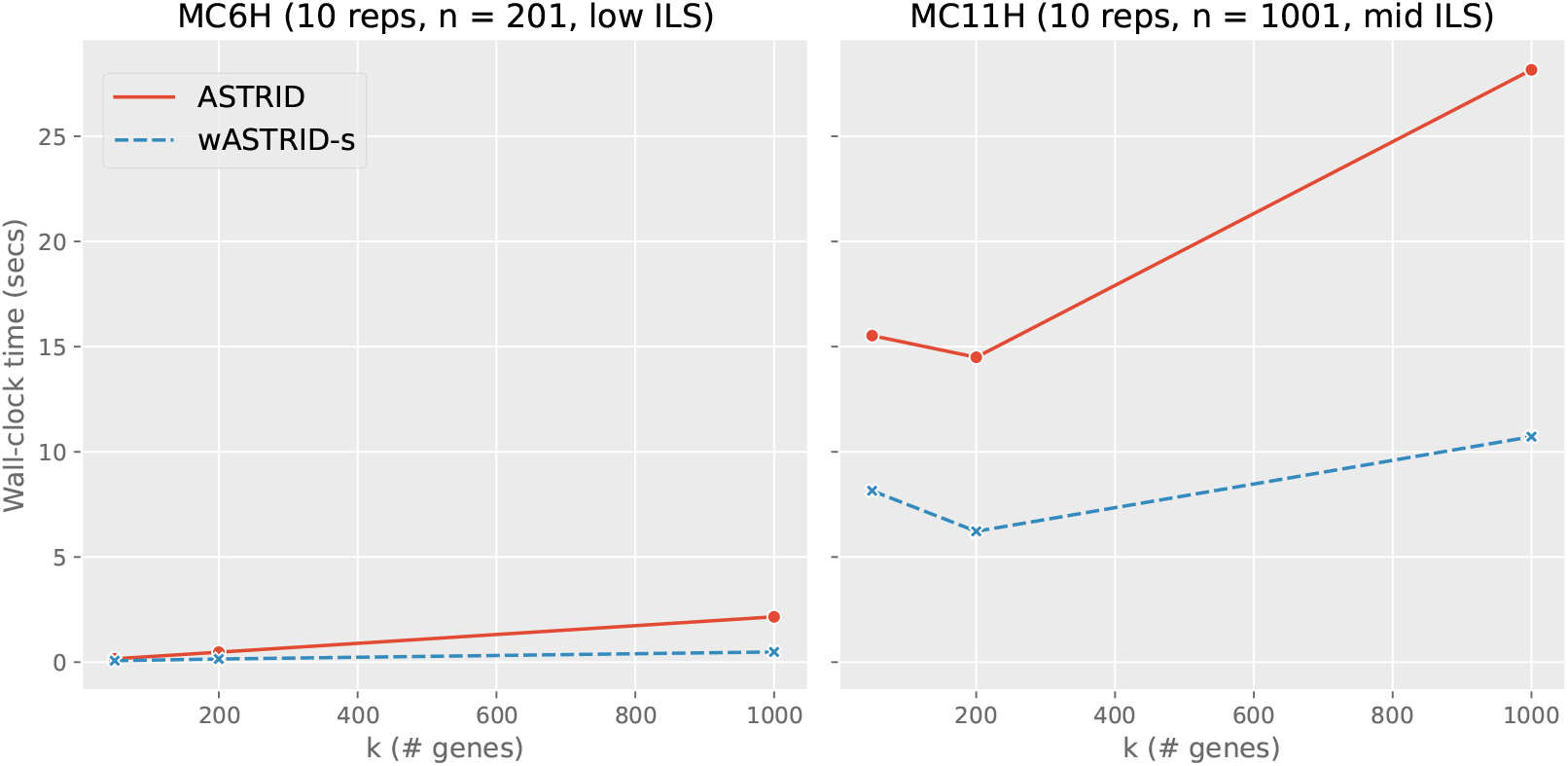
Benchmark of ASTRID vs. wASTRID-s on two datasets comparing the wall-clock running time. Shown data points are averages of ten replicates with *k* (number of genes) ranging in 50, 200, 1000. All methods are run on the same machine with a Intel(R) Xeon(R) CPU (E5-2670 0 @ 2.60GHz).

